# The main duct of von Ebner’s glands is a source of Sox10^+^ taste bud progenitors and susceptible to pathogen infections

**DOI:** 10.1101/2024.05.14.594215

**Authors:** Wenxin Yu, Maria Eleni Kastriti, Mohamed Ishan, Saurav Kumar Choudhary, Naomi Kramer, Md Mamunur Rashid, Hy Gia Truong Do, Zhonghou Wang, Ting Xu, Robert F Schwabe, Kaixiong Ye, Igor Adameyko, Hong-Xiang Liu

**Author notes:** equal contribution to the work (Maria Eleni Kastriti, Mohamed Ishan, Saurav Kumar Choudhary). Correspondence should be sent to Hong-Xiang Liu.

## Abstract

We have recently demonstrated that *Sox10*-expressing (*Sox10*^+^) cells give rise to mainly type-III neuronal taste bud cells that are responsible for sour and salt taste. The two tissue compartments containing *Sox10*^+^ cells in the surrounding of taste buds include the connective tissue core of taste papillae and von Ebner’s glands (vEGs) that are connected to the trench of circumvallate and foliate papillae. In this study, we used inducible Cre mouse models to map the cell lineages of connective tissue (including stromal and Schwann cells) and vEGs and performed single cell RNA-sequencing of the epithelium of *Sox10-Cre/tdT* mouse circumvallate/vEG complex. *In vivo* lineage mapping showed that the distribution of traced cells in circumvallate taste buds was closely linked with that in the vEGs, but not in the connective tissue. *Sox10*, but not the known stem cells marker *Lgr5*, expression was enriched in the cell clusters of main ducts of vEGs that contained abundant proliferating cells, while *Sox10-Cre/tdT* expression was enriched in type-III taste bud cells and excretory ductal cells. Moreover, multiple genes encoding pathogen receptors are enriched in the vEG main ducts. Our data indicate that the main duct of vEGs is a source of *Sox10*^+^ taste bud progenitors and susceptible to pathogen infections.

## Introduction

Taste bud cells have a short lifespan and need to be constantly replaced [1–5]. Thus, stem/progenitor cells in the surrounding tissue compartments are essential for taste bud homeostasis. It has been well documented that the Krt5/14^+^ [6], P63^+^ [6], Sox2^+^ [6–12], Lgr5^+^ [13–15], and *Gli1*^+^ [16] basal cells in taste bud-surrounding lingual epithelium are taste bud progenitors. We have recently found that cells expressing SRY-related HMG-box gene 10 (*Sox10*^+^) give rise to taste bud cells that are mainly type-III with low levels of Krt8 [17]. Using *in situ* hybridization for *Sox10*, we identified two tissue compartments under lingual epithelium that contain *Sox10*^+^ cells, i.e., connective tissue core and von Ebner’s minor salivary glands (vEGs) that are connected to the trench of circumvallate and foliate papillae [17]. These tissues may represent a novel source of progenitors for taste buds.

In our recent study to map the lineage of neural crest exclusively, we used *Sox10-iCreER/tdT^Tmx@E7.5^* and found that traced cells were present in underlying connective tissue but not in taste buds [18], which indicates that taste bud cells do not originate from neural crest and the derived connective tissue. However, the questions remain where and what cells in non-neural crest-derived connective tissue and/or *Sox10*^+^ vEGs originated from endoderm [19–21] are progenitor cells for taste buds.

In this study, we used single cell transcriptomic analyses on the epithelium of circumvallate/vEG complex and multiple inducible Cre mouse models to map the lineage of cells surrounding circumvallate taste buds including the stromal cells and Schwann cells in the connective tissue core of papilla and the cells in vEGs. Our data indicates that *Sox10*^+^ cells in the main duct of vEGs, but not in the connective tissue core of taste papillae, serve as progenitors for taste buds without being transited to basal cells in the stratified tongue epithelium. Moreover, vEG main ducts enrich multiple genes encoding pathogen receptors indicating its susceptibility to pathogen infections.

## Results

### *Sox10-iCreER^T2^/tdT*^Tmx@E7.5^ marks the majority, but not all, of stromal cells in the connective tissue core of circumvallate papilla

The stromal cells found in the mesenchyme and connective tissue of the tongue are predominantly derived from the neural crest, evidenced at both embryonic and postnatal stages [17, 18, 22–24]. However, it is unclear how many non-neural crest-derived cells are present in the connective tissue core of taste papillae. Our recently published data shows that a single-dose tamoxifen to a pregnant dam carrying E7.5 *Sox10-iCreER^T2^/tdT* embryos can effectively mark cells in the lamina propria of the mouse tongue at 8 week of stage [18] when taste buds are mostly mature. In the present study, quantification of the connective tissue cells in the core of circumvallate papilla in adult *Sox10-iCreER^T2^/tdT*^Tmx@E7.5^ mice (Figure 1A, B, and D) revealed that the percentage of *Sox10-iCreER^T2^/tdT*-traced cells versus total cells in the defined area of connective tissue was 79.94±3.39% (X±SD). Of note, tdT^+^ cells were neither found in the ducts (Figure 1B) and acini (Suppl Figure 1) of von Ebner’s glands (vEGs), nor in circumvallate taste buds marked by Krt8 immunosignals (Figure 1B). Moreover, tdT^+^ cells were not observed in the taste bud-surrounding basal tongue epithelial cells (Figure 1B) that are known taste bud progenitors [3, 6, 25–29].

**Figure 1.**
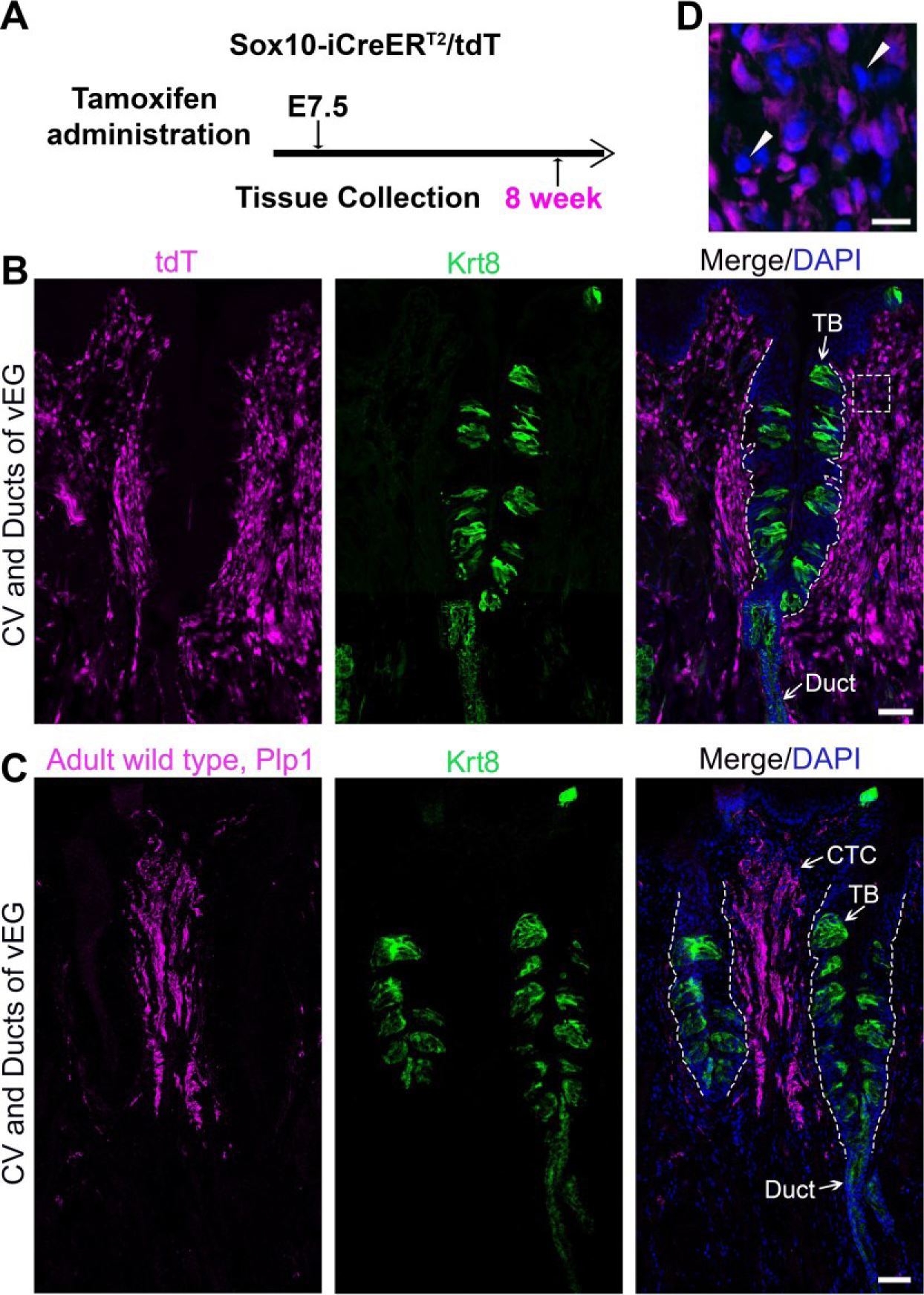
The distribution of neural crest-derived cells and Plp1-expressing (Plp1^+^) cells in the connective tissue core of circumvallate papilla. **<u>A</u>:** A schematic diagram to illustrate the experimental design using *Sox10-iCreER^T2^/tdT* mice for neural crest cell mapping in B. **<u>B and C</u>:** Single-plane laser scanning confocal photomicrographs of coronal sections of circumvallate papilla to demonstrate the distribution of *Sox10-iCreER^T2^/tdT^+^*-traced (B) and Plp1^+^ (C) cells (magenta). Arrowheads point to Krt8^+^ immunostained (green) cells in taste buds (TB), ductal (Duct) cells of von Ebner’s gland, and connective tissue core (CTC) of circumvallate papilla. Arrowheads point to the tdT^-^ cells. White dashed lines mark the borders between the epithelium and underlying connective tissue. Scale bars: 50 μm in B and C; 10 μm in the high-power image from the squared area in B.

This data suggests that up to 20% of the connective tissue cells were not traced by *Sox10-iCreER^T2^/tdT*^Tmx@E7.5^ and are potentially non-neural crest derived. This data keeps the possibility of *Sox10*^+^ connective tissue cells as taste bud progenitors an open question warranting further studies.

### *Plp1-CreER^T^*, *Plp1-CreER^T2^*, and *Vimentin-CreER* maps cells extensively in the lamina propria, but not in circumvallate taste buds nor von Ebner’s glands (vEGs)

In the connective tissue core, we observed two abundant and largely distinct subpopulations of cells that express either Proteolipid protein 1 (*Plp1*) (Figure 1C), an intrinsic membrane protein specifically expressed in Schwann cells in the peripheral nervous system [30], or Vimentin [17, 18, 22–24], an intermediate filament protein widely used as a marker for mesenchymal stromal cells [31]. To verify the potential contribution of connective tissue cells to taste buds, we used inducible Cre mouse lines in which CreER is under the control of the promoter of *Plp1* or *Vimentin*.

In the P11 *Plp1-CreER^T^/tdT*^Tmx@P1-10d^ (Figure 2A-B), *Plp1-CreER^T2^/YFP*^Tmx@P1-10d^ (Figure 2A, 2C), and *Vimentin-CreER/tdT*^Tmx@P1-10d^ (Figure 3A, 3D) mice, tdT^+^/YFP^+^ cells were abundantly distributed in the connective tissue core of circumvallate papilla after daily tamoxifen treatments from P1 to P10. No tdT^+^/YFP^+^ cells were found in taste buds and vEGs (Figure 2B-D; Suppl Figure 2A-B, 2D). This data demonstrated that our tamoxifen treatment paradigm was sufficient to induce DNA recombination for tracing Plp1^+^ and Vimentin^+^ cell lineages in the connective tissue. The absence of tdT^+^/YFP^+^ cells in taste buds at this early time point makes these mouse models eligible for taste bud cell derivation assay at later time points.

**Figure 2.**
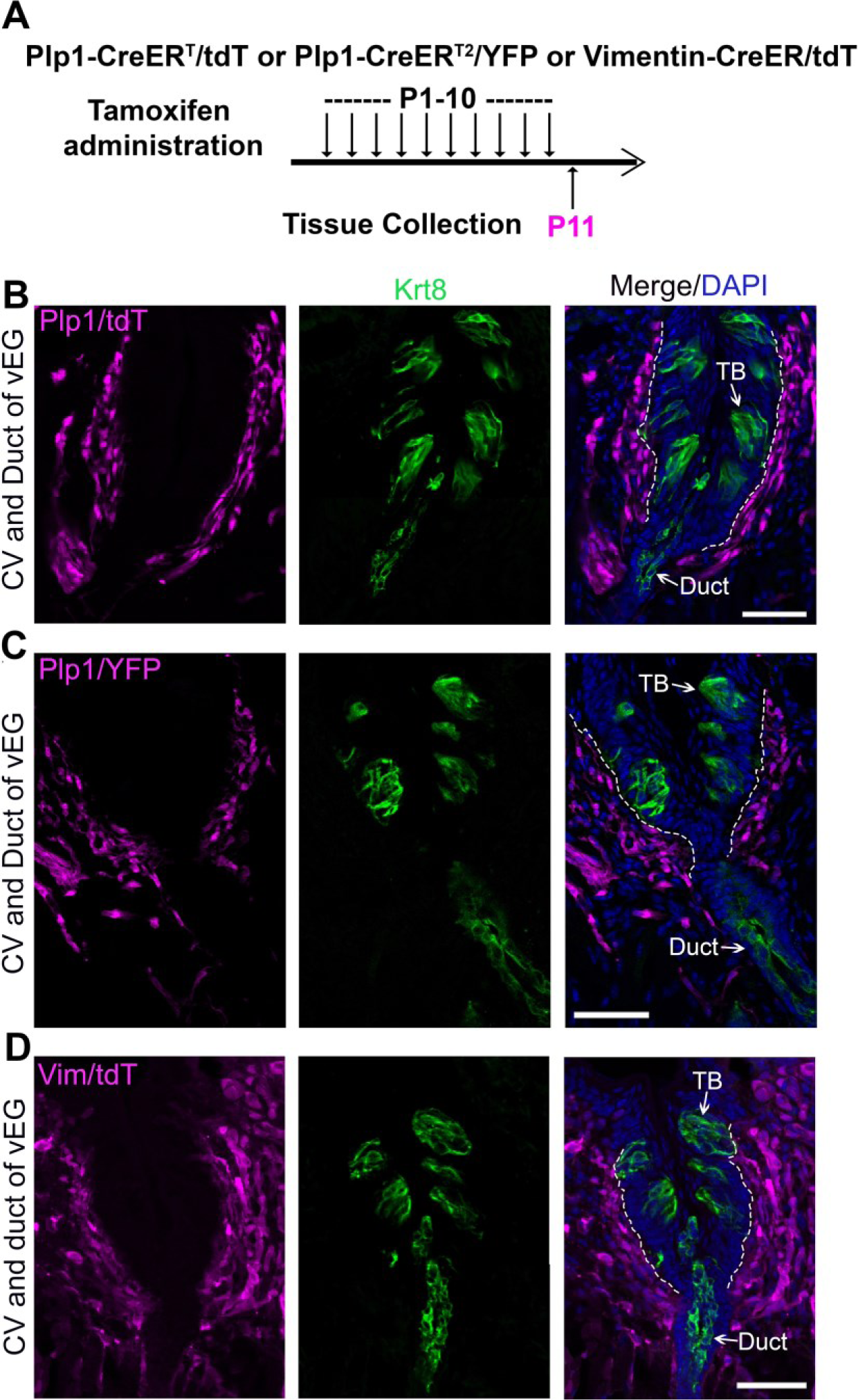
Tamoxifen administration from P1 to P10 efficiently induce labeling of connective tissue cells in the circumvallate papilla using *Plp1-CreER^T^/tdT* or *YFP* mice (Schwann cells) and *Vimentin-CreER/tdT* mice (stromal cells). **<u>A</u>**: A schematic diagram to illustrate the timeline of daily tamoxifen administration from P1-10 and tissue collection at P11. **<u>B-D</u>**: Single-plane laser scanning confocal images of circumvallate papilla and von Ebner’s gland on coronal sections of *Plp1-CreER^T^/tdT* (B), *Plp1-CreER/YFP* (C) and *Vimentin-CreER/tdT* (D) mice. Traced cells, seen as tdT^+^ (B and D) and YFP^+^ (C) cells (magenta), are abundant in the connective tissue core of circumvallate papilla. Arrows point to Krt8^+^ immunostained (green) cells in taste buds (TB), ductal (Duct) cells of von Ebner’s gland. White dashed lines mark the borders between the epithelium and surrounding connective tissue. Scale bars: 50 μm in all images.

**Figure 3.**
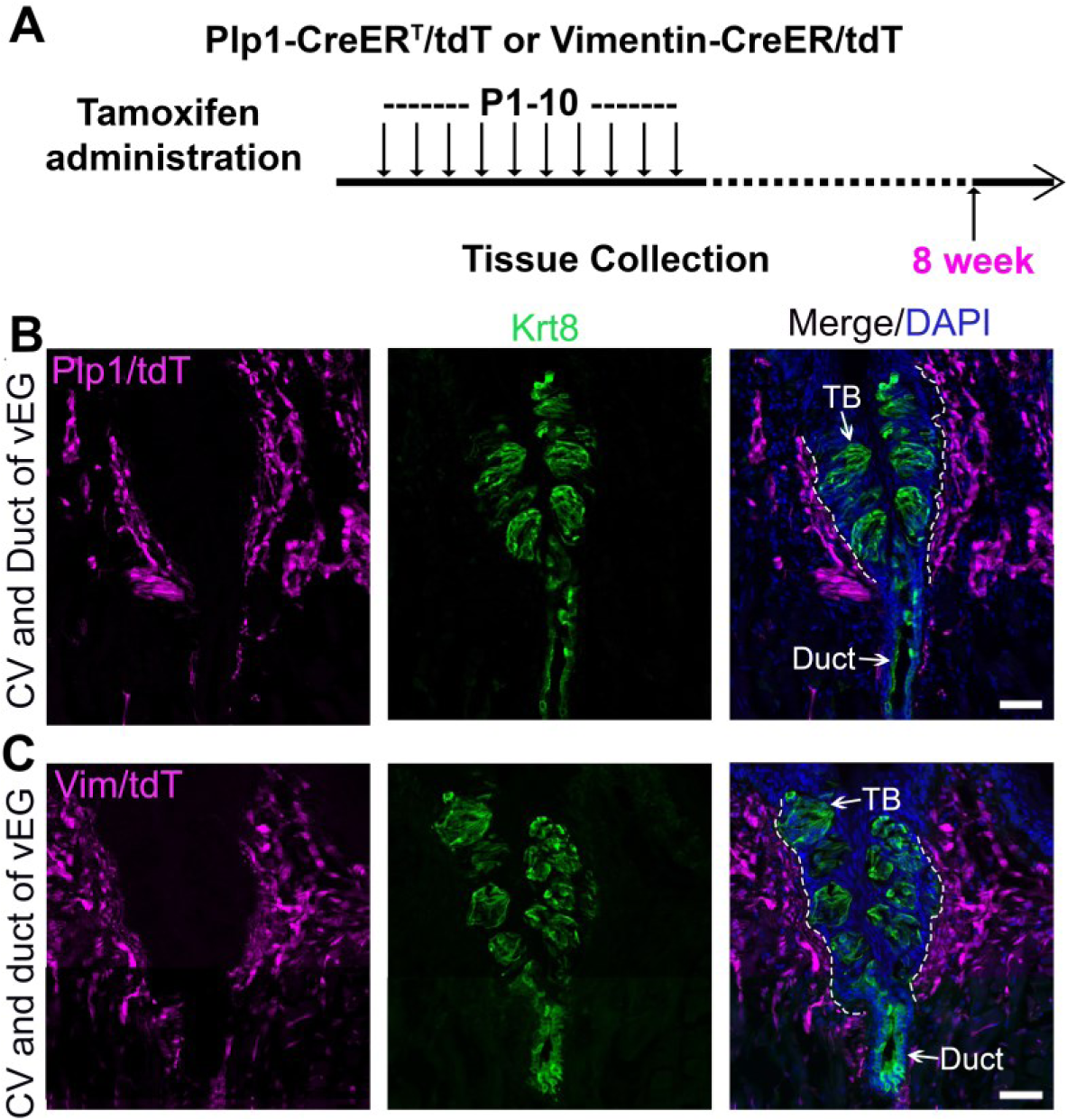
*Plp1-CreER^T^/tdT*-traced Schwann cells and *Vimentin-CreER/tdT*-traced stromal cells do not contribute to circumvallate taste buds. **<u>A</u>**: A schematic diagram to illustrate the timeline of daily tamoxifen administration from P1-10 and tissue collection at 8 weeks of age. **<u>B-D</u>**: Single-plane laser scanning confocal images of circumvallate papilla and von Ebner’s gland on coronal sections of *Plp1-CreER^T^/tdT* (B) and and *Vimentin-CreER/tdT* (C) mice. tdT^+^ (magenta) are present in the connective tissue core but not in taste buds of circumvallate papilla. Arrows point to Krt8^+^ immunostained (green) cells in taste buds (TB), ductal (Duct) cells of von Ebner’s gland. White dashed lines mark the borders between the epithelium and surrounding connective tissue. Scale bars: 50 μm for all images.

Given that type-III taste bud cells derived mainly from *Sox10*^+^ progenitors [17] have a half-life of 22 days [4] vs. ∼8-12 days on average for all taste bud cells [1, 32], a prolonged time window was given after the termination of tamoxifen treatments and 8-week-old mice were harvested for examination after ten daily doses of tamoxifen from P1 to P10 (Figure 3A). Both *Plp1-CreER^T^/tdT* and *Vimentin-CreER/tdT* mouse models showed sustained extensive distribution of tdT^+^ cells in the connective tissue core of the circumvallate papilla (Figure 3B-C). A thorough examination of serial sections revealed that no tdT^+^ cells were observed in any of the taste buds (Figure 3B-C). Of note, tdT^+^ cells were also absent in the vEG ducts (Figure 3B-C) and acini (Suppl Figure 2A, 2C, 2E). In addition, taste bud-surrounding tongue epithelium was free from tdT tracing signals in *Plp1-CreERT/tdT* and *Vimentin-CreER/tdT* mice at all stages examined (Figure 2B-D; Figure 3B-C).

### Postnatal induction of *Sox10-iCreER^T2^/tdT* and *Sox10-CreER^T2^/YFP* concurrently marks von Ebner’s glands and circumvallate taste buds

Ruling out the derivation of taste buds from *Sox10*^+^ cells in the neural crest and connective tissue, we evaluated the distribution of *Sox10*^+^ cell lineage in circumvallate taste buds and its association with that in vEGs. Using *Sox10-iCreER^T2^/tdT* and *Sox10-CreER^T2^/YFP* mouse models with daily tamoxifen administration from P1 to P10 (Figure 4A; Suppl Figure 3A), we observed that at P11, vEGs (Figure 4B, 4D; Suppl Figure 3B) and connective tissue (Figure 4B, 4D) were clearly labeled in abundance. It is important to note that at this early time point tdT^+^/YFP^+^ cells were not seen in circumvallate taste buds nor the taste bud-surrounding lingual epithelial cells including the basal cells (Figure 4B, 4D). When given a period of approximately 7 weeks after the termination of tamoxifen treatments, tdT^+^ and YFP^+^ cells were seen, albeit infrequently, in circumvallate taste buds (Figure 4C, 4E, 4F-G) in both *Sox10-iCreER^T2^/tdT*^Tmx@P1-10d^ and *Sox10-CreER^T2^/YFP*^Tmx@P1-10d^ mice (Figure 4A; Suppl Figure 3A). Concurrently, tdT^+^ cells sustained in vEGs (Figure 4C; Suppl Figure 3C).

**Figure 4.**
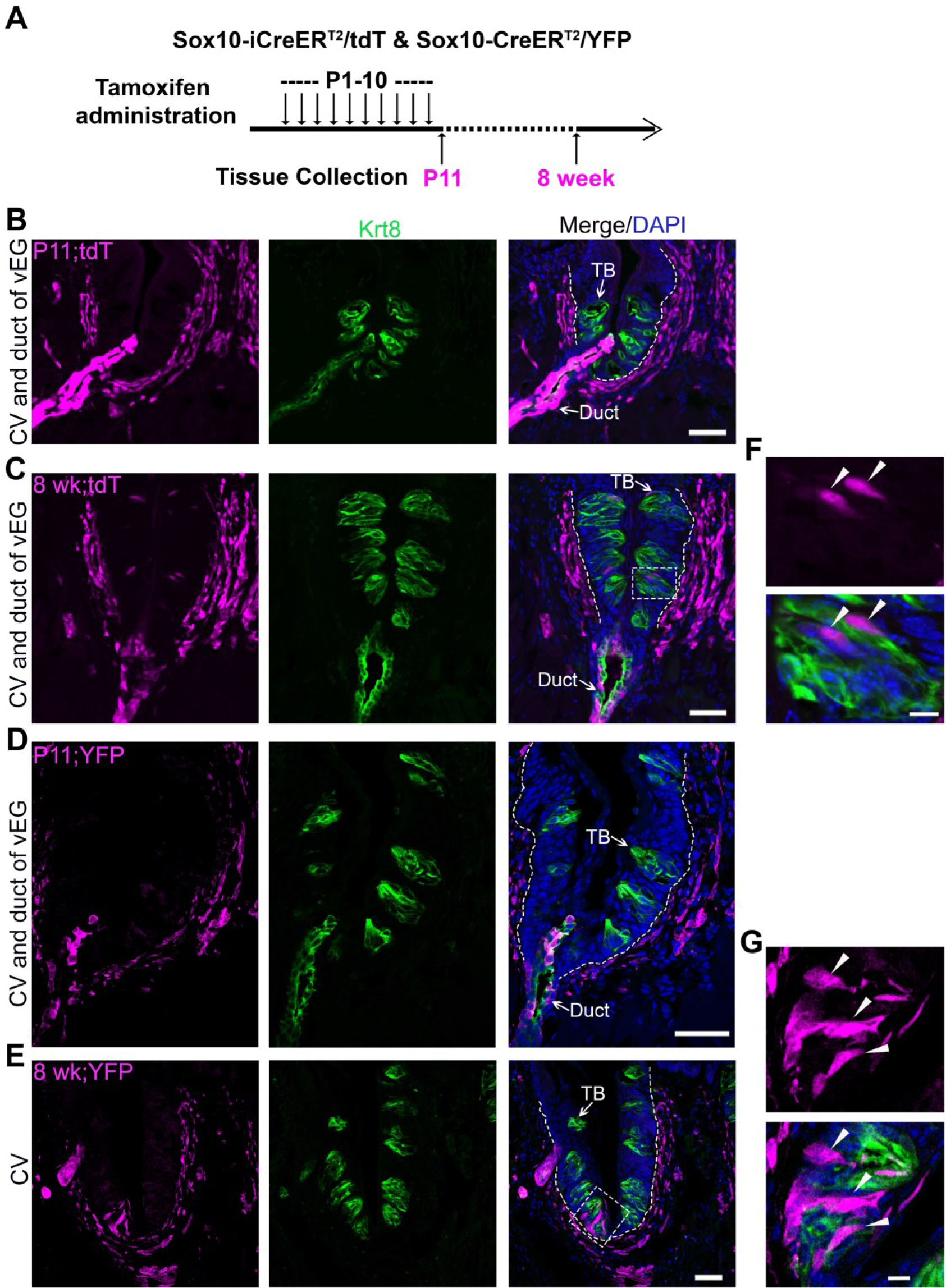
Cell labeling and lineage mapping of *Sox*10^+^ cells in Von Ebner’s glands and circumvallate taste buds. **<u>A</u>:** A schematic diagram to show the experimental paradigm in *Sox10-iCreER^T2^/tdT* and *Sox10-CreER/YFP* mice. **<u>B-E</u>:** Single-plane laser scanning confocal photomicrographs on coronal sections of circumvallate papilla. The distribution of *Sox10-iCreER^T2^/tdT* (B-C) and *Sox10-CreER/YFP*-traced (D-E) cells (magenta) of P11 (B and D) and 8-week-old (C and E) mice with tamoxifen administration from P1 to P10. **<u>F-G</u>:** High-power images of the area enclosed in the dashed square in C (F) and E (G). Arrows point to Krt8^+^ immunostained (green) cells in taste buds (TB) and ductal (Duct) cells of von Ebner’s gland. Arrowheads in F-G point to the tdT^+^ and YFP^+^ cells in taste buds. White dashed lines mark the borders between the epithelium and surrounding connective tissue. Scale bars: 50 μm in B-E, 10 μm in F-G.

### Long-term consecutive tamoxifen administration results in a more abundant distribution of *Sox10-iCreER^T2^*-traced cells in circumvallate taste buds

To test whether the infrequent *Sox10-iCreER^T2^/tdT*^Tmx@P1-10d^ labeling of taste bud cells was caused by transient Sox10 expression or short lifespan of *Sox10*^+^ progenitors in vEGs, consecutive tamoxifen administration was performed in *Sox10-iCreER^T2^/tdT* mice from P1 day to 8 weeks or up to 16 weeks (Figure 5A; Suppl Figure 4A). Indeed, after 8 weeks of tamoxifen treatment, *Sox10-iCreER^T2^/tdT*-labeled cells were abundantly distributed within circumvallate taste buds (Figure 5B-C) and concurrently in vEGs (Figure 5B; Suppl Figure 4B). The labeled taste bud cells depicted the typical fusiform shape of differentiated taste cells (Figure 5C). A similar labeling pattern was observed in *Sox10-iCreER^T2^/tdT* mice treated with tamoxifen for 16 weeks from birth (Figure 5D-E; Suppl Figure 4C). Of note, *Sox10-iCreER^T2^/tdT*-labeled cells were not observed in taste bud-surrounding lingual epithelium in all stages examined (Figure 5B, 5D).

**Figure 5.**
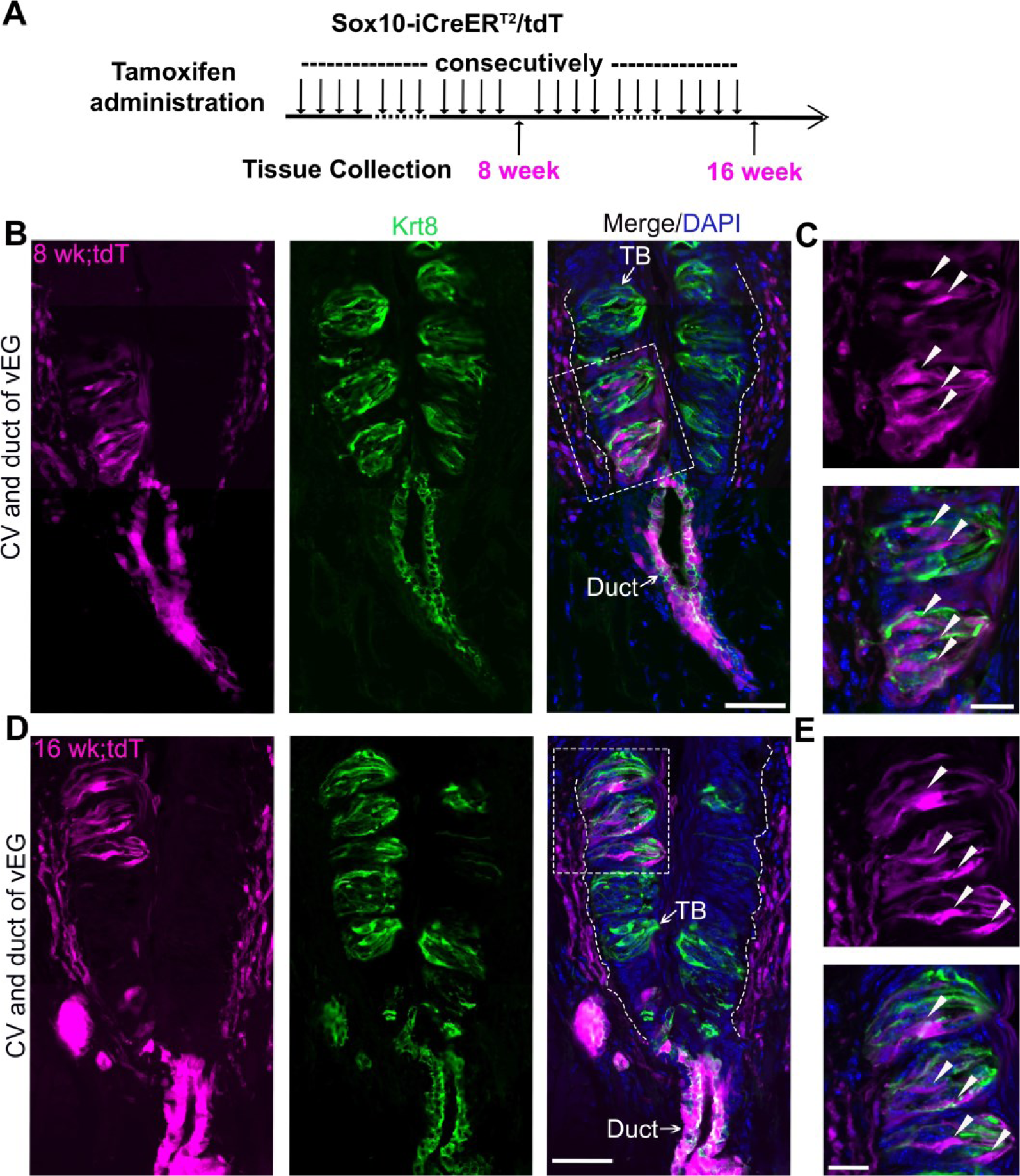
Long-term (8-wk or 16-wk) tracing leads to abundant distribution of *Sox10-iCreER^T2^/tdT*-labeled cells in circumvallate taste buds concurrently with those in Von Ebner’s glands. **<u>A</u>:** A schematic diagram to illustrate the experimental paradigm for tamoxifen administration and tissue collection from *Sox10-iCreER^T2^/tdT* mice at 8 or 16 weeks. **<u>B-E</u>:** Single-plane laser scanning confocal images of a coronal circumvallate papilla section at 8-week (B-C) or 16-week (D-E). tdT^+^ cells (magenta) were concurrently distributed in circumvallate taste buds and von Ebner’s glands in addition to that in connective tissue. C and E are high-power images from the area enclosed in the dashed square shown in B and D respectively. Arrows point to Krt8^+^ immunostained (green) cells in taste buds (TB) and ductal (Duct) cells of von Ebner’s gland. Arrowheads in C and E point to the tdT^+^ cells in taste buds. White dashed lines mark the borders between the epithelium and surrounding connective tissue. Scale bars: 50 μm in B and D; 20 μm in C and E.

### Single cell transcriptomic analyses reveal *Sox10* expression in vEG main ducts and *Sox10-Cre/tdT* expression in Type-III taste cells

To identify the epithelial cell population of *Sox10^+^* cells and derived cells in the epithelium of circumvallate papilla/vEG complex (Fig. 6A_1_), single cell RNA-sequencing (scRNA-Seq) was performed using dissociated cells from the epithelial sheets of *Sox10-Cre/tdT* mice (Figure 6A_2-3_). *Sox10-Cre/tdT*-traced signals were robust in the vEG ducts that are connected to the trench of circumvallate papilla (Figure 6A_3_). At least 13 cell clusters were identified according to their associated signature genes (Figure 6B). Type I-III taste bud cells were distinctly identifiable. Cell types existing in the stratified epithelium were found including the basal, supra-basal and superficial (corneocytes) epithelial layers. Three clusters of glandular cells (main duct, excretory duct, acinar) were detected. In addition, a few other small clusters were identified representing erythrocytes, immune cells, and stromal cells.

**Figure 6.**
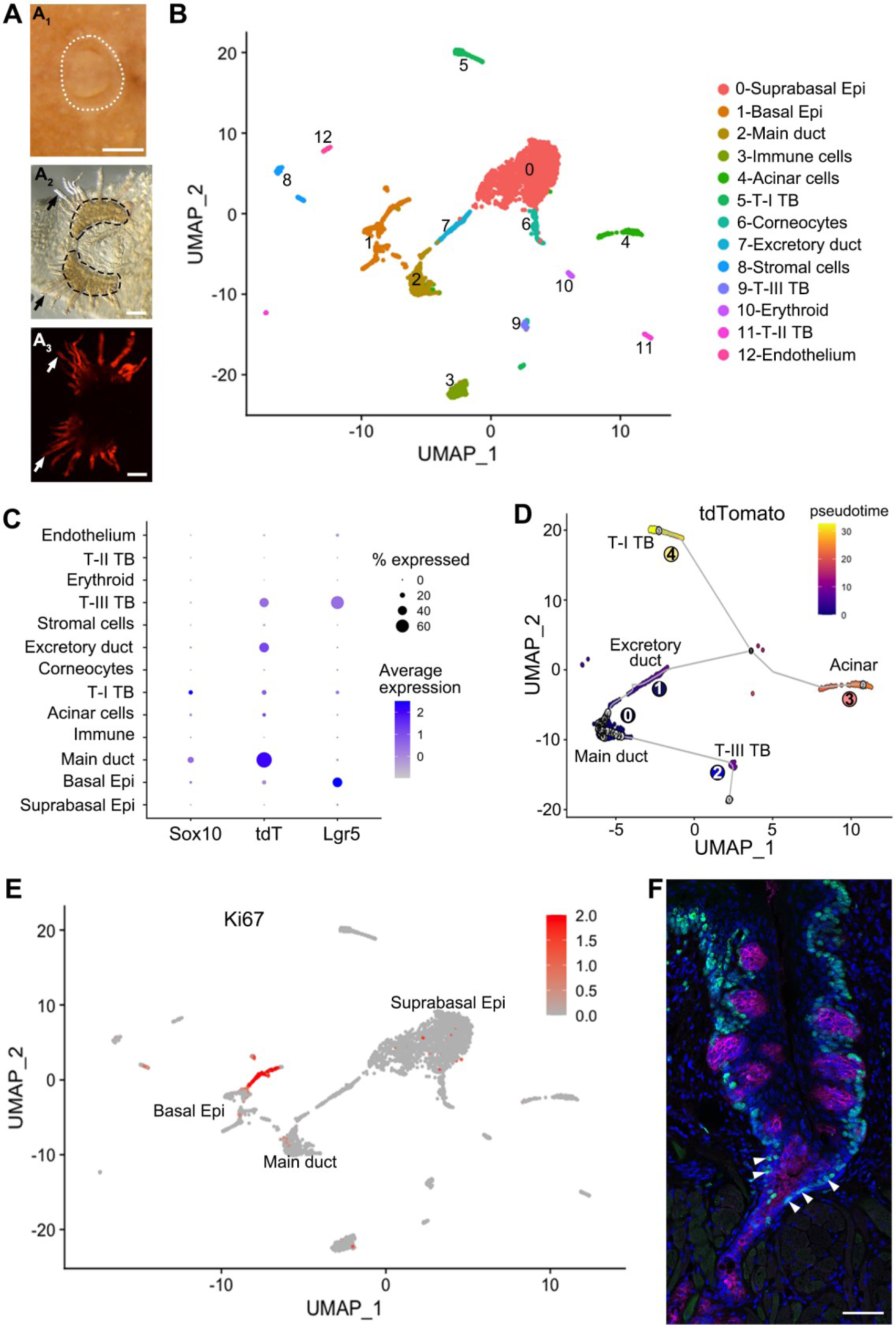
Sox10 expression is enriched in the main duct of vEG while Sox10-Cre/tdT marks type-III taste bud cells and excretory ducts. **<u>A</u>:** whole mount images showing the circumvallate papilla and peeled epithelial sheet of circumvallate/vEG complex for cell dissociation. White dotted lines (A_1_) encircle the circumvallate papilla in postnatal tongue. Black dashed lines (A_2_) encircle the trenches of circumvallate papilla. Dark arrows (A_2_) and white arrows (A_3_) point to the ducts of vEG respectively. Epi: epithelium; T-I, T-II, T-III: type-I, II, III; TB: taste bud. **<u>B</u>:** t-SNE map to illustrate the 13 cell clusters identified in the scRNA-seq data from target tissues shown in A. **<u>C</u>:** Dot plot to illustrate gene expression of *Sox10, tdT, Ki67* across cell clusters. **<u>D</u>:** Pseudotime analysis on the t-SNE map by setting the illustrate the cell position along a trajectory. **<u>E</u>:** Feature plots of Ki67 transcripts on the UMAP showing cell cluster expressing Ki67. **<u>F</u>:** Single-plane laser scanning confocal photomicrographs of coronal sections of circumvallate papilla to demonstrate the distribution of Ki67^+^ cells (green). White Arrowheads point to Ki67^+^ immunostained cells in the transition area between trench of circumvallate papilla and main duct of vEG. Taste buds are marked by Krt8 immunosignals (magenta). Scale bars: 250 μm in A_1_; 100 μm in A_2_, A_3_; 50 μm in F.

Among these cell clusters, *Sox10* expression was enriched in the main duct of vEGs and a small proportion of differentiated type-I taste bud cells (Figure 6C). In contrast, tdT expression marking the *Sox10*^+^ cell lineage was enriched in type-III taste bud cells and excretory vEG ductal cells in which *Sox10* expression was not detected. Moreover, tdT expression was detected at a high level in over 60% of the main ductal cells. Of note, Lgr5, the well-known taste bud stem cell marker, was expressed in basal epithelial cells (e.g., known progenitors for taste buds) and type-III taste bud cells, but not detected in any types of vEG cells.

To infer the differentiation trajectory of *Sox10-Cre/tdT*^+^ cells, pseudotime analysis indicated ordering of cell clusters from the vEG main duct to the two immediate neighbor cell compartments, i.e., excretory duct and taste bud (mainly type-III) cells (Figure 6D). The two clusters of further order of cells include acinar and type-I taste bud cells that express *Sox10* in themselves.

To identify the proliferating and potential progenitors for taste buds, enrichment of Ki67 expression was found in basal and supra-basal epithelial cells, as well as in the main ducts of vEGs (Figure 6E). Immunosignals of Ki67 confirmed the abundant distribution of Ki67^+^ proliferating cells in the main ducts of vEGs in addition to the basal and some supra-basal epithelial cells surrounding taste buds (6F).

### The main ducts of vEGs express receptors for multiple types of pathogens

We have reported the distinct tropism of multiple viruses in the anterior tongue epithelial cells [33]. In this study, the posterior distribution of viral receptor expression was performed to understand the vulnerability of different cell types in the circumvallate papilla/vEG epithelium to infectious diseases (Fig. 7). We found that different pathogen receptors were enriched in distinct cell clusters. Across all 13 identified cell clusters, the main ducts of vEGs have an overall high frequency in expressing key viral entry factors for SARS-CoV-2 (*Tmprss2*), HCoV-229E (*Anpep*), influenza virus (*St3gal4*), and MERS-CoV45 (*Dpp4* and *Tnfrsf1a*), and multiple receptors involved in the innate immune system. Similarly, the excretory ducts of vEGs enrich multiple pathogen receptors. In contrast, basal tongue epithelial cells (i.e., the well-known taste bud progenitor) depicted a low frequency and abundance of pathogen receptor gene expression. Differentiated taste bud cells showed distinct and relatively low frequency of expression of pathogen receptors.

**Figure 7.**
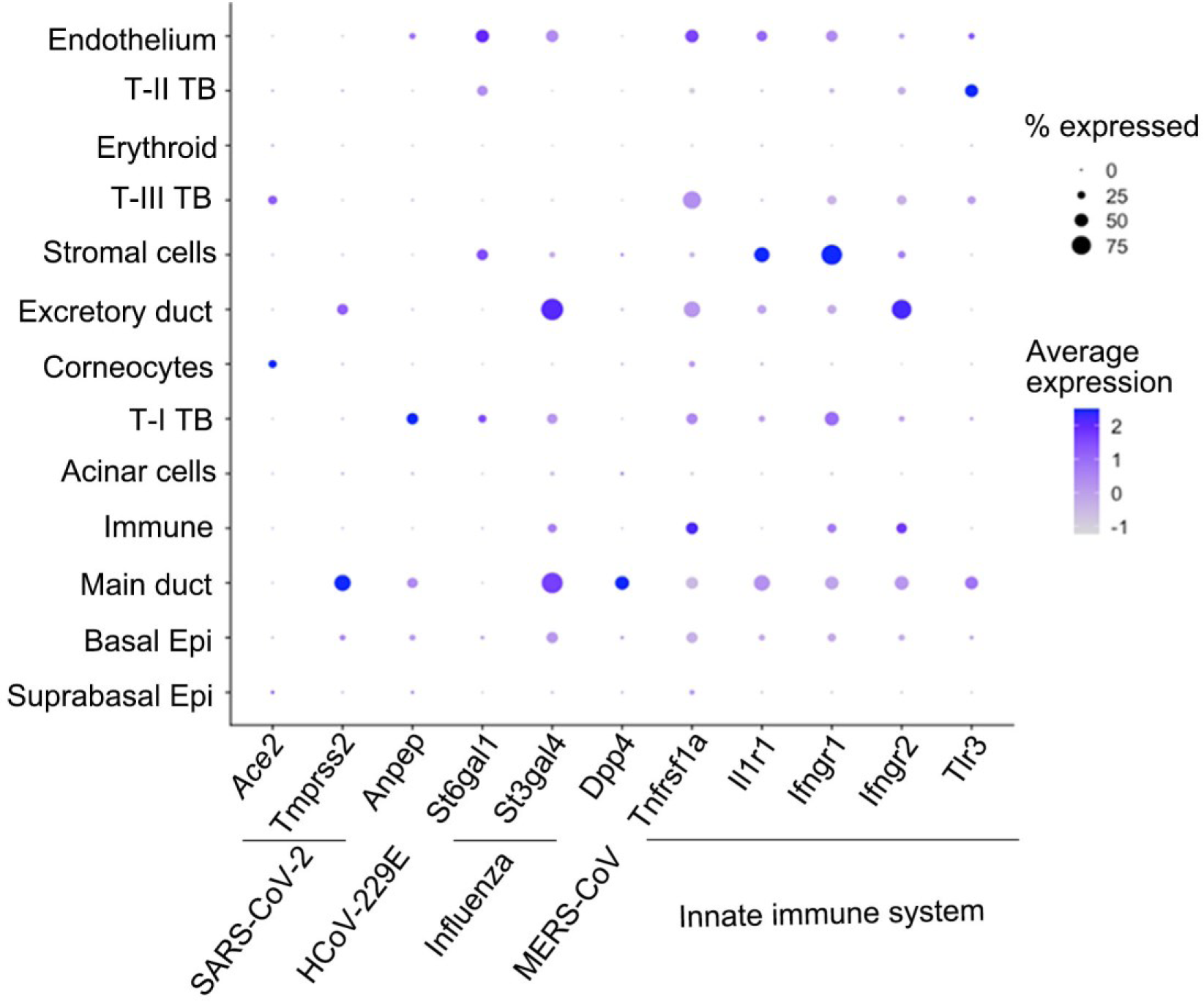
Dot plot to illustrate gene expression (log(UMIs count/ 10 000 + 1)) across the 13 cell clusters in the scRNA-seq data set. Genes related to SARS-CoV-2, HCoV-229E, influenza, MERS-CoV, and the innate immune system are included. The size of the dots represents the proportion of gene-expressing cells, and the color intensity of the dots represents the average level of the gene expression.

## Discussion

### *Sox10*^+^ cells in von Ebner’s glands (vEGs), but not those in connective tissue, serve as progenitors for circumvallate taste buds

Our recent studies have shown that cells expressing *Sox10* (*Sox10*^+^) are the progenitors of type-III neuronal taste cells responsible for sour and salt taste [17]. Three tissue compartments that contain *Sox10*^+^ cells include the neural crest in mouse embryos, connective tissue core of taste papillae and vEGs in postnatal mice [17]. Specific mapping of neural crest cell lineage by administering a solitary dose of tamoxifen to *Sox10-iCreER^T2^/tdT*^Tmx@E7.5^ mice resulted in marking of up to 80% of connective tissue cells (neural crest-derived) in the core of the circumvallate papilla. However, this did not trace taste bud cells [18]. As a result, it remains uncertain whether *Sox10*^+^ connective tissue cells are the progenitors of taste buds. Further lineage mapping of the two abundant cell subpopulations (i.e., *Plp1*^+^ Schwann and *Vimentin*^+^ stromal cells) in the connective tissue core of taste papillae indicates that the derived cells are not distributed in taste buds. The observations in our previous study using *Vimentin-CreER*-driven membrane-bound GFP reporter [34], in which mGFP signals were seen in taste buds [24], may be attributed to the extended nerve fiber or fibroblast lamellipodia from connective tissue.

After confirming that taste buds do not originate from connective tissue cells, the only possible candidate left for *Sox10*^+^ taste bud progenitors is vEGs -- the minor salivary glands that are connected to the bottom of the trenches of circumvallate and foliate papillae in the posterior tongue [35, 36]. Indeed, using inducible CreER mouse models under the control of the *Sox10* promoter to trace cells in distinct tissue compartments, the distribution of traced cells in circumvallate taste buds is tightly associated with that in vEGs, but not with that in the connective tissue core of circumvallate papilla.

Of note, considering the importance of proper use of CreER and Cre reporter mouse lines, the concurrent distribution of traced cells in vEGs with those in circumvallate taste buds was confirmed with two different Cre reporters and two independently developed CreER under the control of *Sox10* promoter (*Sox10-iCreER^T2^/tdT* and *Sox10-CreER^T2^/YFP*). Together with the observations that taste buds exist in the circular structure of tissue transplants in the deep layer under the circumvallate papilla trench [37], our data demonstrate that vEGs are the source of *Sox10*^+^ progenitors for taste buds (Figure 8).

**Figure 8.**
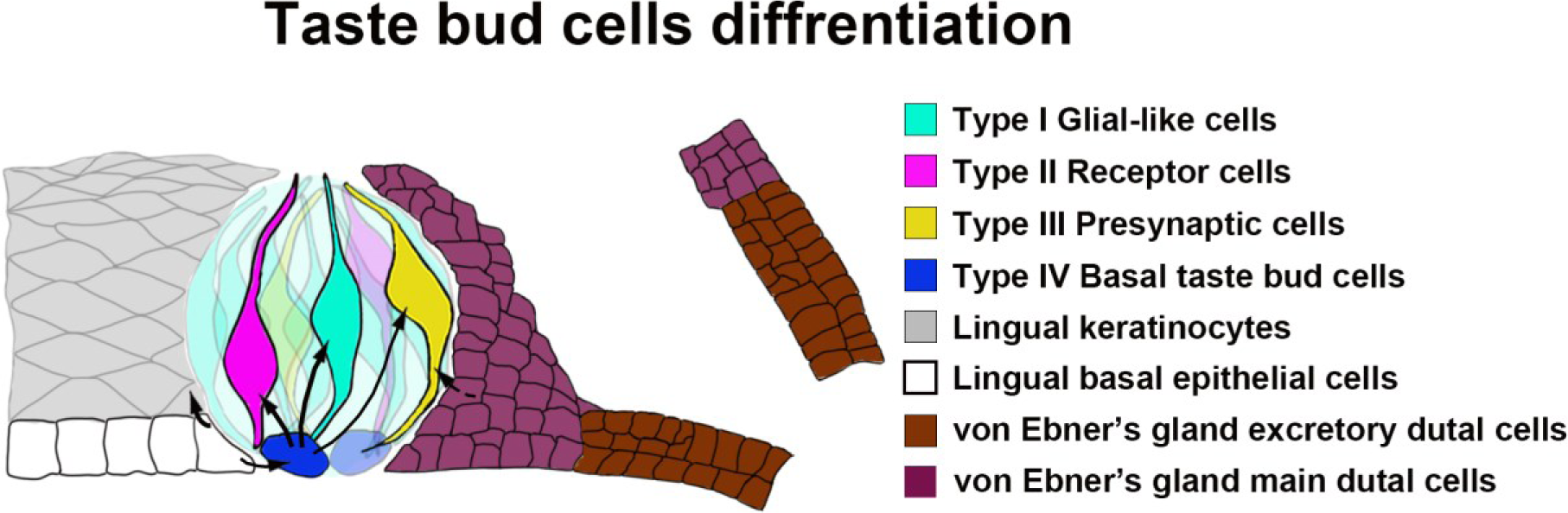
A schematic diagram illustrating the tissue compartments contributing to circumvallate taste buds. Our data indicate that Sox10-expressing cells in von Ebner’s gland serve as stem/progenitor cells for taste bud cells in the circumvallate papilla.

### The main ducts of vEGs are the source of Sox10^+^ taste bud progenitors susceptible to pathogen infections

The vEGs are one type of the minor salivary glands that are connected and open to the trough of circumvallate and foliate taste papillae [38]. Minor salivary glands are structurally comprised of the endpieces called acini and the ducts including interlobular, intercalated, excretory and collecting (main) ducts. In mice, the vEGs at vallate papilla are located under circumvallate taste buds and composed of serous acini and interlobular excretory ducts [39]. At the opening of vEG duct (main duct) the columnar epithelium transitions to stratified squamous epithelium [38, 39].

Our scRNA-seq data in circumvallate papilla/vEG complex from *Sox10-Cre/tdT* mice allows us to identify the heterogeneity of epithelial cell population and to compare the *Sox10*^+^ and tdT^+^ traced cells at the single cell resolution. The enrichment of *Sox10* expression in vEG main duct contrasts that of *tdT^+^* signals (marking the *Sox10^+^* cell lineage) in the main duct and additional cells in its immediate neighbor tissues, i.e., type-III taste bud cells and excretory duct, which provides a clear indication that vEG main ducts give rise to excretory ducts and type-III taste bud cells. This is also supported by the cell trajectory analysis and abundant distribution of proliferating cells in the main ducts of vEGs.

Regarding the derivation of other types of taste cells, both *Sox10* and *tdT* expression was enriched in the same small percentage of type-I cells suggesting that the *Sox10-Cre/tdT*-traced signals [17] were due to the *Sox10* expression in the type-I cells. In this study, we did not detect *tdT* expression in type-II taste cells. A large-scale scRNA-Seq study may be needed to identify a low number of traced cells, if any type-II taste cells.

*Lgr5*, the known taste progenitor marker for taste buds [13–15], was recently reported to be expressed in the vEGs [40]. In contrast, our scRNA-Seq data revealed the expression of Lgr5 in basal tongue epithelial cells but not in the vEGs. Likely the Lgr5-traced vEGs are derived from the basal tongue epithelial cells as it has been reported that the circumvallate trench gives rise to vEGs [40]. Thus, *Sox10*^+^ taste progenitors in main duct of vEG are a distinct population of taste progenitors from those marked by Lgr5. Further studies are needed to identify the specific ductal cell type(s) that migrate and differentiate to taste bud cells.

While traced cells are present in both circumvallate taste buds and vEGs, no traced cells are found in the lingual epithelium surrounding taste buds. This suggests that the ductal cells of Ebner’s glands migrate and differentiate directly into taste bud cells, rather than being “transited” by becoming the taste bud-surrounding lingual epithelial cells. It has been recently reported that *Nkx2-2*^+^ taste cells are committed to the type-III lineage [41]. *Sox10*^+^ vEG ductal progenitors may initiate the expression of *Nkx2-2* for the terminal differentiation to type-III taste cells. Overall, our data add vEG main ducts to the progenitor sources for circumvallate taste buds (Figure 8).

### vEG ductal taste bud progenitors express *Sox10* transiently and are susceptible to pathogen infections

When comparing the distribution of both *Sox10-iCreER^T2^/tdT* and *Sox10-CreER^T2^/YFP*-traced cells in vEGs with those in circumvallate taste buds, our data reveal the contribution dynamics of *Sox10*^+^ vEGs progenitors to taste buds. After inducing Cre recombination in neonatal mice for 10 days, *Sox10-CreER* efficiently labeled vEGs, but only a limited number of cells were traced in the circumvallate taste buds several weeks later. A prolonged series of tamoxifen treatments is required to mark more taste bud cells, suggesting a transient *Sox10* expression or a short lifespan of *Sox10*^+^ vEG taste bud progenitors.

In our previous study [33], scRNA-Seq profiling of anterior mouse tongue epithelial cells [42] revealed that molecules for various viruses’ entries into cells are enriched in different cell clusters. Our current scRNA-Seq data from the circumvallate/vEG complex reveals distinct tropism of multiple viruses in the posterior tongue region in a different pattern from that in the anterior tongue. Combined data in both anterior and posterior tongue epithelium provide insights into the mechanisms underlying taste loss in different infectious diseases. For example, detection of SARS-CoV-2 receptor gene *Ace2* in taste cells (type-III) suggests that direct SARS-CoV-2 infection of taste cells may be the cause of acute taste loss. The expression of co-receptor *Tmprss2* in vEG main ducts and *Ace2* in the taste bud-surrounding epithelial cells may cause a sequela, i.e., long-term taste dysfunction. As the newly found taste bud progenitor source, vEG main ducts enrich receptor genes encoding multiple pathogens making the organ susceptible to infections.

### Derivation of connective tissue cells in the tongue

The tongue is a complex and highly organized organ in which the mesenchymal/connective tissue cells serve as scaffolds to compartmentalize different tissues [43, 44]. In the connective tissue core of taste papillae, we observed two profuse and largely distinct subpopulations of cells that express Vimentin, an intermediate filament protein widely used as a marker for mesenchymal stromal cells [31] and Proteolipid protein 1 (Plp1), an intrinsic membrane protein specifically expressed in Schwann cells in the peripheral nervous system [30]. The cells in the tongue mesenchyme/connective tissue are largely derived from the cranial neural crest [17, 18, 22–24, 45, 46]. Multiple mouse models have been used to map neural crest lineages in the tongue, including *Wnt1-Cre* [47–54], *P0-Cre* [23], *Dermo1-Cre* [24, 55, 56], and *Sox10-Cre* [57, 58]. The Cre models label cells in the tongue mesenchyme of embryos and connective tissue in the postnatal mouse tongue [17, 24]. However, none of these models labels neural crest cells exclusively or labels all neural crest cells [59, 60]. There is no available data regarding the proportion of neural crest- and non-neural crest-derived cells.

*Sox10* is specifically expressed in the neural crest in early embryos [17, 61]. Inducible *Sox10-CreER* [18] with tamoxifen administration at E7.5 [62, 63] is optimal in mapping neural crest cell lineages exclusively. A single-dose tamoxifen administration to the pregnant dam carrying E7.5 *Sox10-iCreER^T2^/tdT* embryos results in extensive labeling of tongue mesenchymal cells at E12.5 [18]. In the present study, quantification of *Sox10-iCreER^T2^/tdT* traced cells in the connective tissue core of circumvallate papilla indicates that at least 80% of the cells are neural-crest derived. An increase in tamoxifen dose(s) may lead to a higher proportion of labeled cells; however, low embryo viability limited the study. To confirm if up to 20% of connective tissue cells in the tongue originate from mesoderm, specific mouse models need to be analyzed.

Together with the previous findings [18, 37], our data in the present study supports the concept that *Sox10*^+^ cells in the main ducts of vEGs can migrate and differentiate to circumvallate taste bud cells that are mainly type-III for sour and salt (Figure 8). These *Sox10*^+^ cells represent an additional source of taste bud progenitors in distinction from the basal lingual epithelial cells. The differentiation of Ebner’s gland cells into taste bud cells for transducing gustatory signals represents a novel format of tissue-tissue interactions in the circumvallate/Ebner’s gland complex in addition to gland’s secreting saliva to facilitate taste perception.

## Materials and methods

### Animals

Mice were used in this study (Table 1). The animal use was approved by the Institutional Animal Care and Use Committee at The University of Georgia and the Medical University of Vienna and complied with the National Institutes of Health Guidelines for the care and use of animals for research.

**Table 1.**
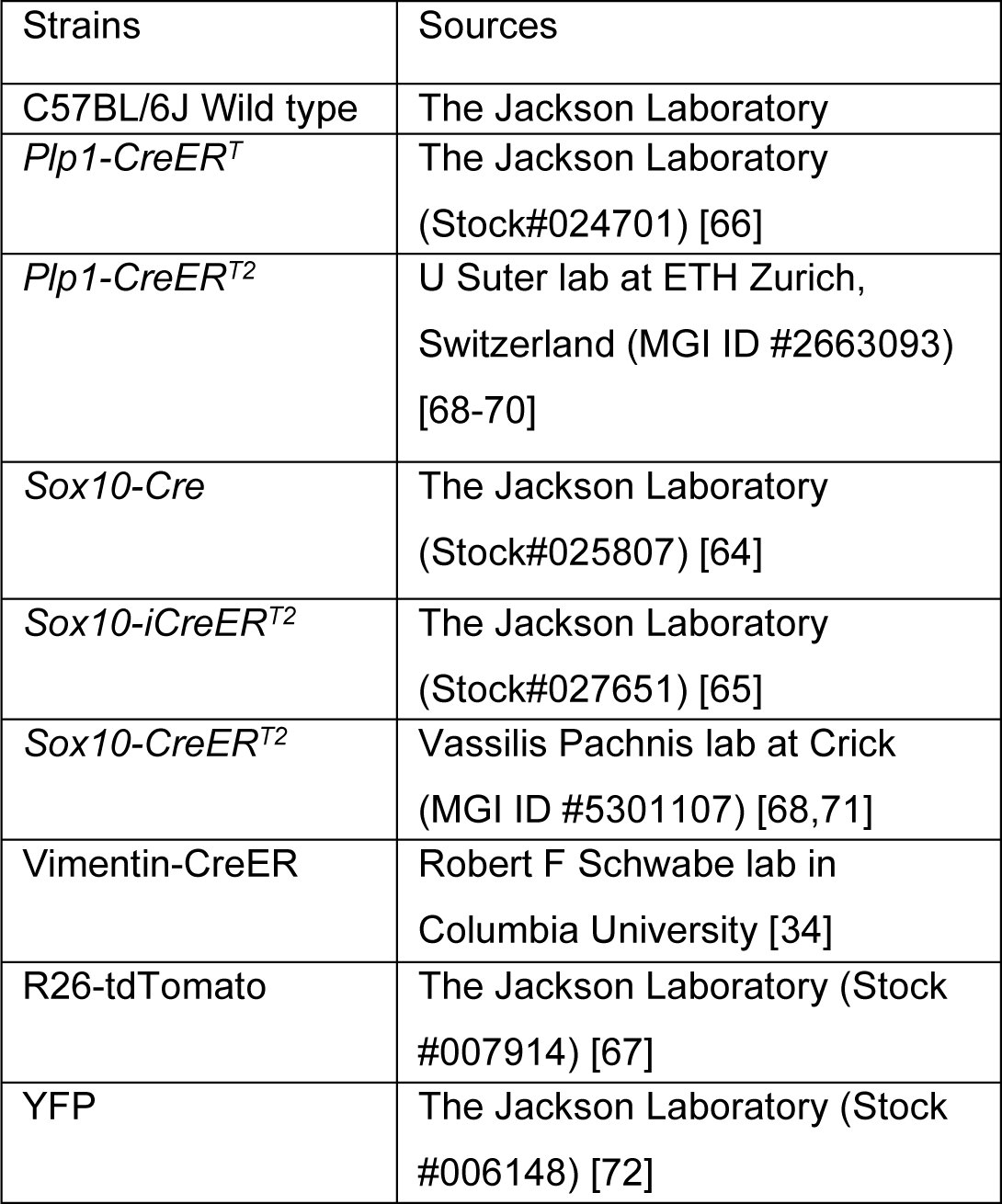
Mouse strains that were used.

| Strains | Sources |
| --- | --- |
| C57BL/6J Wild type | The Jackson Laboratory |
| *Plp1-CreER<sup>T</sup>* | The Jackson Laboratory<br>(Stock#024701) [66] |
| *Plp1-CreER<sup>T2</sup>* | U Suter lab at ETH Zurich,<br>Switzerland (MGI ID #2663093)<br>[68-70] |
| *Sox10-Cre* | The Jackson Laboratory<br>(Stock#025807) [64] |
| *Sox10-iCreER<sup>T2</sup>* | The Jackson Laboratory<br>(Stock#027651) [65] |
| *Sox10-CreER<sup>T2</sup>* | Vassilis Pachnis lab at Crick<br>(MGI ID #5301107) [68,71] |
| Vimentin-CreER | Robert F Schwabe lab in<br>Columbia University [34] |
| R26-tdTomato | The Jackson Laboratory (Stock<br>#007914) [67] |
| YFP | The Jackson Laboratory (Stock<br>#006148) [72] |

Mice were maintained in the animal facility of the Animal and Dairy Science department at the University of Georgia at 22℃ under 12-hr day/night cycles. *Sox10-Cre* [64], *Sox10-iCreER^T2^* [65], *Plp1-CreER^T^* [66], and *Vimentin-CreER* [34] mice were bred with *R26-tdTomato* (hereafter tdT) Cre reporter mice [67] respectively to generate *Sox10-Cre/tdT, Sox10-iCreER^T2^/tdT*, *Plp1-CreER^T^/tdT*, and *Vimentin-CreER/tdT* mice. Being aware of the importance of use of proper CreER and reporter mouse lines, two independent CreER (*Plp1-CreER^T2^* [68–70] and *Sox10-CreER^T2^* [68, 71]) mouse lines crossed with a different Cre reporter *YFP* (yellow fluorescent protein) [72] were used to verify our observations. The posterior tongues containing circumvallate taste buds and von Ebner’s glands were provided by Dr. Igor Adameyko’s lab at the Medical University of Vienna. Data from female and male mice did not show significant differences regarding the distribution of labeled cells in each strain; therefore, females and males were grouped for analyses. FVB/NJ wild-type mice (The Jackson Laboratory, Stock#001800) were used to breed in parallel for fostering cesarean-born *Sox10-iCreER^T2^/tdT* pups. Polymerase chain reaction (PCR) genotyping was performed using the following primers listed in Table 2.

**Table 2.**
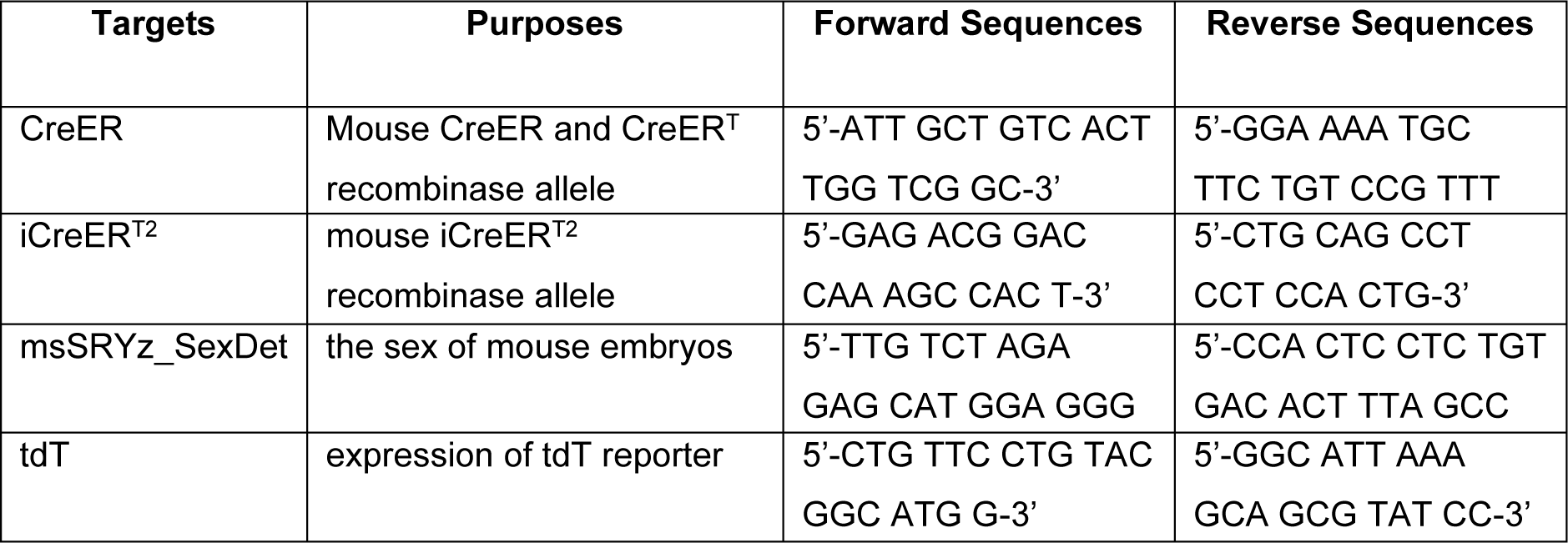
PCR genotyping primers.

### Tamoxifen administration and fostering of pups

Tamoxifen (Tmx) (Cat No. T5648; Sigma-Aldrich, Inc, St. Louis, MO) was dissolved in the mixture of ethanol and corn oil (1:9) at the working concentration of 10.0 or 11.1 mg/mL. CreER-mediated DNA recombination was induced by oral gavage of 0.1 mL tamoxifen solution to the pregnant or nursing dams. Vehicle controls were treated with corn oil in the dams or littermates with the same genotype in parallel.

For labeling neural crest cell lineages, timed pregnant mice were used. Noon of the day of vaginal plug detection in mice was designated as embryonic day (E) 0.5. To induce Cre recombination in neural crest cells, a single dose of 0.1 mL tamoxifen solution was given to the pregnant dams that carried E7.5 *Sox10-iCreER^T2^/tdT* embryos through oral gavage using 16 G x 38 mm polyurethane feeding tubes (Cat No. FTPU-16-50, Instech Laboratories, Inc, Plymouth Meeting, PA). To resolve the dystocia problem caused by tamoxifen treatment in pregnant mice, cesarean sections were performed to deliver the *Sox10-iCreER^T2^/tdT* embryos at E18.5. The delivered pups were fostered by FVB/J nursing dams immediately after cesarean birth.

To activate CreER-induced recombination in neonatal mice, nursing dams were given 0.1 mL tamoxifen daily from birth (P1) to P10. The nursing dams for *Sox10-CreER^T2^/YFP* pups received 0.1 mL tamoxifen every other day. For mice with prolonged tamoxifen administration, 0.1 mL tamoxifen solution was given to the nursing dams once every three days until weaning on day 28, followed by oral gavage to weaned mice once every three days until 8 weeks or 16 weeks.

### Tissue collection

Postnatal (P) mice were harvested on day 11, week 8 or 16 of age (n=3 for both vehicles- and tamoxifen-treated at each stage). Mice were euthanized with CO_2_ followed by cervical dislocation. For scRNA-Seq, the tongue was immediately dissected for cell dissociation as previously described [73]. For immunohistochemistry on tissue sections, transcardial perfusion was performed using 10 mL warm 0.1 M PBS, 10 mL warm, and 20 mL cold 2% PFA in 0.1 M PBS. Tongues were dissected and further fixed in 2% PFA at 4℃ for 2 hr.

### Single cell RNA-sequencing (scRNA-Seq)

*SOX10-Cre/tdT* mice were used at 8 week of age when the proportion of traced cells in circumvallate taste buds is abundant and plateaued [17]. The separation of the epithelium of circumvallate papilla region and cell dissociation were performed as previously described [73]. A total of 8 circumvallate/vEG epithelial sheets were pooled for single cell isolation. Dissociated cells (for a target of sequencing ∼10,000 cells) of circumvallate papilla/vEG epithelium were submitted to the Univ of Georgia – Georgia Genomics and Bioinformatics Core for scRNA-seq library preparation (10X Genomics) and Illumina sequencing.

The reads from sequencing were processed using Cell Ranger (v7.0.0) for alignment, filtering, barcode counting, UMI counting and cell counting. A customized reference genome was built by adding gene tdTomato (tdT) to mouse reference genome (mm10). Analysis of scRNA-seq dataset was performed with R package Seurat (v5.0.1). We followed the analysis pipeline recommended by Satija Lab (https://satijalab.org). In cell quality control, cells with less than 500 total RNA counts and less than 250 detected features were excluded. Additionally, cells with mitochondrial gene exceeding 20% were filtered out to remove potentially stressed or dying cells. Data was normalized and PCAs were calculated using “RunPCA”, Uniform Manifold Approximation and Projection (UMAP) analyses have been done using “RunUMAP” (dims=1:20). “FindNeighbours” was used for defining the distance between the cells and later these were grouped using “FindClusters” with resolution 0.1. Subsequently, clusters were annotated using marker genes from the literature. For trajectory analysis, Monocle3 (v1.3.4) was employed, following the recommended analysis pipeline by Trapnell Lab (https://cole-trapnell-lab.github.io/monocle3/) [74–76].

### Immunohistochemistry

Fixed tongues were cryoprotected in 30% sucrose in 0.1 M PBS for at least 48 hr at 4℃. The circumvallate papilla was dissected from the tongues and trimmed under a stereomicroscope, embedded in O.C.T. (Cat. No. 23-730-571, Thermo Fisher Scientific, Pittsburgh, PA), and rapidly frozen. Coronal sections were cut at 8 μm in thickness and mounted onto charged slides (Fisher brand™ Superfrost™ Plus Microscope Slides, Cat. No. 12-550-15, Thermo Fisher Scientific, Pittsburgh, PA). Non-specific binding was blocked with 10% normal donkey serum (Cat. No. SLBW2097, Sigma-Aldrich, Inc, St. Louis, MO) in 0.1 M PBS containing 0.3% Triton X-100 (Cat. No. X100-100ML, Sigma-Aldrich, Inc, St. Louis, MO) for 30 min, followed by overnight incubation with primary antibody against GFP (1:500, SKU: GFP-1020, Aves Labs), or Keratin (Krt) 8 (1:500, TROMA-1, Developmental Studies Hybridoma Bank), or Ki67 (1:200, ab16667, ABCAM, MA), or Plp1 (1:500, #28702, Cell Signaling Technology, Danvers, MA) diluted with 0.1 M PBS containing 0.3% Triton X-100 and 1% normal donkey serum. After rinses in 0.1 M PBS (3 times, 10 min each), sections were incubated with Alexa Fluor® 647 (Krt8) secondary antibody (1:500, Invitrogen, Eugene, OR) and Alexa Fluor® 488 (Ki67, Plp1) secondary antibody (1:500, Invitrogen, Eugene, OR) for 1 hr at room temperature. Sections were rinsed and then counterstained with DAPI (200 ng/mL, Cat. No. D1306, Thermo Fisher Scientific, Pittsburgh, PA) for 10 min at room temperature. After rinses with 0.1 M PBS followed by dipping in Milli-Q water (DirectQ® 3 UV water purification system, Millipore, MA). Sections were air-dried and cover slipped with Prolong® diamond antifade mounting medium (Cat. No. P36970, Thermo Fisher Scientific, Pittsburgh, PA).

### Photomicroscopy and quantifications

Immunostained sections were thoroughly examined under a fluorescent light microscope (EVOS FL, Thermo Fisher Scientific, Pittsburgh, PA) and images were taken using a laser scanning confocal microscope in the UGA Microscopy Core (Zeiss LSM 710, Zeiss, Germany). The total cells and *Sox10-iCreER^T2^/tdT*-unlabeled cells in connective tissue immediately under circumvallate taste buds were quantified on single-plane confocal images from 8-week-old *Sox10-iCreER^T2^/tdT*^Tmx@E8^ mice (n=3) using Photoshop software. The region of underlying connective tissue within 100 μm in distance to the basement membrane of the circumvallate papilla epithelium was included for the quantification. For each mouse, three sections were selected for analysis, i.e., the section with the deepest trench and biggest area of circumvallate papilla, and those two anterior and posterior sides of sections at 40 μm away from the section with the deepest trench. Only the cells with a clear DAPI^+^ nucleus were counted. A cell with a clear DAPI^+^ nucleus and without any tdT signals in the cell was regarded as a *Sox10-iCreER^T2^/tdT* non-labeled cell. The proportion of neural crest-derived connective tissue cells was calculated using the formula of [DAPI^+^ tdT^+^ cells / DAPI^+^ (tdT^-^ + tdT^+^) cells].

## Acknowledgments

This study was supported by the National Institutes of Health, grant number R21DC018910 and R21DC018089 to HXL, and by ERC Synergy Grant (856529 - KILL - OR-DIFFERENTIAT - ERC-2019-SyG), Paradifference Foundation, Knut and Alice Wallenberg Foundation, Austrian Science Fund, Swedish Research Council, Hjärnfonden (FO2023-0255) to IA. The English was edited by Francisca Gibson Burnley (Regenerative Bioscience Center, Department of Animal and Dairy Science, University of Georgia).

## Author contributions

WY and HXL conceived of the idea, designed the experiment, wrote the manuscript. WY, MEK, MI, MMR, NK, ZW, TX and HXL performed the experiments. WY, SKC, HGTD, NK and HXL processed and analyzed the data. MEK, RFS, KY, and IA provided research materials. KY, IA and HXL supervised the work.

