## Supplemental Figures 1-4 for "The main duct of von Ebner’s glands is a source of Sox10^+^ taste bud progenitors and susceptible to pathogen infections"

[#] equal contribution to the work.

Supplemental Figure 1

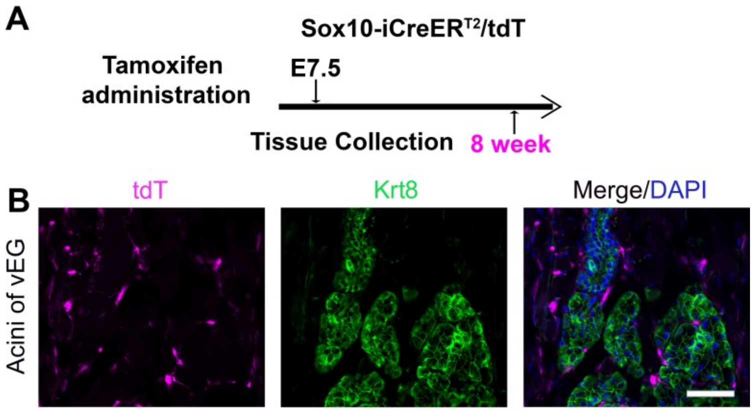


**Supplemental Figure 1.** Neural crest cell lineage mapping in the acini of von Ebner's glands**. A:** A schematic diagram to illustrate the experimental design using *Sox10-iCreER*$^{T2}$*/tdT* mice for neural crest cell mapping. **B:** Single-plane laser scanning confocal photomicrographs of von Ebner's glands on coronal sections. *Sox10-iCreER*$^{T2}$*/tdT*-labeled cells (magenta) are scattered in the surrounding tissue but not seen in the glands. Scale bars: 50 μm.

Supplemental Figure 2

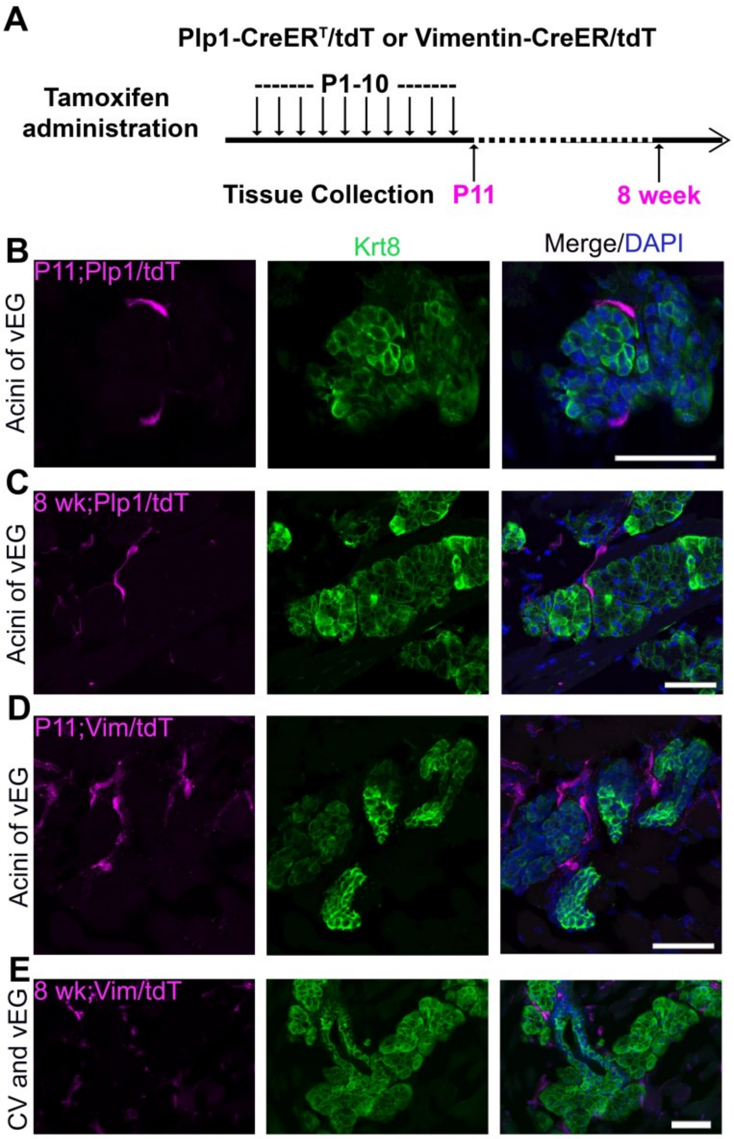


**Supplemental Figure 2.** Cell labeling and lineage mapping in von Ebner's gland using *Plp1-CreER$^T$/tdT* and *Vimentin-CreER/tdT* mice with tamoxifen administration from P1 to P10. **A**: A schematic diagram to illustrate the timeline of daily tamoxifen administration from P1-10 and tissue collection at P11 day and 8 weeks. **B-E**: Single-plane laser scanning confocal images of von Ebner's glands on coronal sections of circumvallate papilla region in *Plp1-CreER$^T$/tdT* (B-C) and *Vimentin-CreER/tdT* (D-E) mice. Scale bars: 50 μm in all images.

Supplemental Figure 3

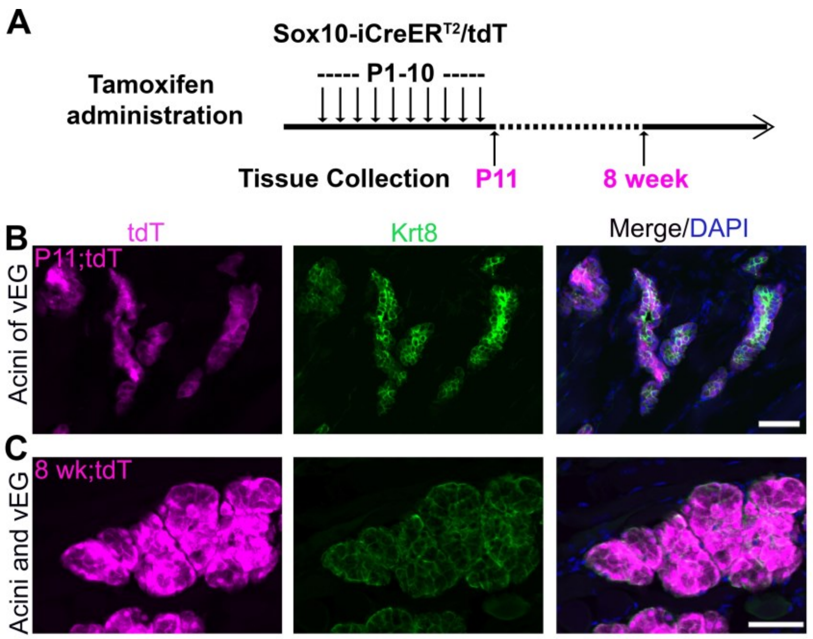


**Supplemental Figure 3.** Cell labeling and lineage mapping of $Sox10^{+}$ cells in Von Ebner's glands. **A:** A schematic diagram to show the experimental paradigm in *Sox10-iCreER$^{T2}$/tdT* mice. **B-C:** Single-plane laser scanning confocal photomicrographs on coronal sections of von Ebner's gland. Following tamoxifen administration from P1 to P10, Sox10-iCreER$^{T2}$/tdT$^{+}$ cells (magenta) are abundantly distributed in the ducts and acini of von Ebner's glands at P11 (B) and 8- weeks (C) of mice. Scale bars: 50 μm in all images.

Supplemental Figure 4

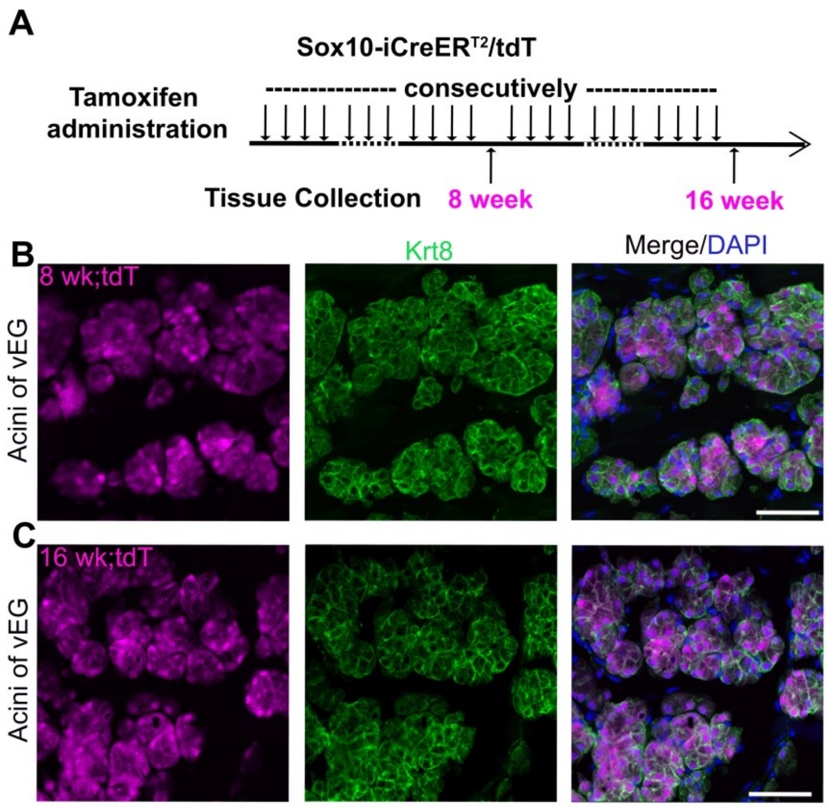


**Supplemental Figure 4.** Abundant distribution of *Sox10-iCreER$^{T2}$/tdT*-labeled cells in acini of Von Ebner's glands in mice treated with tamoxifen for a long-term (8-wk or 16-wk) . **A:** A schematic diagram to illustrate the experimental paradigm for tamoxifen administration and tissue collection from *Sox10-iCreER$^{T2}$/tdT* mice at 8 or 16 weeks. **B-C:** Single-plane laser scanning confocal images of von Ebner's glands on coronal sections of circumvallate papilla region in 8-week-old (B) and 16-week-old (C) mice. *Sox10-iCreER$^{T2}$/tdT$^{+}$* cells (magenta) are abundantly distributed in the acini of von Ebner's glands. Scale bars: 50 µm in all images.
